# A hidden biomass flow - 10,000 tons of insects fly over Switzerland

**DOI:** 10.64898/2026.03.02.708718

**Authors:** Felix Liechti, Birgen Haest, Silke Bauer

**Affiliations:** Department of Environmental Sciences, Zoology, University of Basel, Basel, Switzerland; Swiss Birdradar Solutions AG, Winterthur, Switzerland; Swiss Ornithological Institute, Sempach, Switzerland; Federal Institute for Forest, Snow and Landscape Research WSL, Birmensdorf, Switzerland; ETH Zürich, Zürich, Switzerland

**Keywords:** insects, biomass, aerial, migration

## Abstract

Flying insects are the most diverse and abundant group of animals using the airspace to travel and spread across regions. Through movements that can span multiple life stages and generations, aerial insect migrations shape the ecosystems where they emerge but also connect ecosystems across sometimes vast distances. Yet even basic data on the abundance of aerial insects is still scarce. Their small sizes and often high-altitude movements make it challenging to describe flyways or measure this transport of biomass within or between regions. We used a remote-sensing approach with vertical looking radars to quantify insect migration across Switzerland. Switzerland harbours complex terrain, with the Alps, Jura mountains, and pre-Alpine lowlands featuring starkly contrasting topography. At three locations within these three main habitats, we recorded a total of 6.6 million medium- to large-sized individual insects during the 8-month study period. Extrapolating this to the whole of Switzerland results in an estimated number of 21 billion insects. Adding the presumed additional proportion of small insects, this translates to a rough estimate of 10,000 tons of insect biomass across Switzerland per year. Our results show that insect migration remains substantial even in topographically complex landscapes and seems strikingly synchronized over a relatively wide area. Mountain passes in particular act as key conduits for insects traversing the Alps, with many crossings occurring at high altitudes and with air temperatures below 10 °C. Nevertheless, our study is only a first step unraveling insect movements across space and time and their large-scale synchronicity.

**Significance Statement:** Insect abundance is of great importance for most terrestrial food chains, including human agriculture. While ground-based studies have elucidated many relationships within trophic systems, very little is known about the effects of their large-scale movements through the airspace. We show how insect movements, dispersal and migration, are spatially and temporarily correlated across a wide area of complex terrain and provide a rough estimate of the insect biomass involved. Temperature is a crucial factor in the extent of activity; however, our results show that low temperatures (<10°C) do not prevent massive insect migrations when crossing mountain ranges. Our results demonstrate the large, coordinated flow of insect biomass across the landscape, which needs to be studied in much more detail to better understand the spread of pests and beneficial insects, their impact on local communities, and the transmission of diseases.

## Introduction

Paste your introduction herMany animals migrate through the air to habitats aligned with specific stages of their life cycle (Nilsson et al., 2025; Reynolds et al., 2014). Among these, flying insects are the most abundant (Chapman et al., 2015). Insects play critical roles in species communities, trophic structures and ecosystem functioning (Bauer et al., 2024) and their services and disservices in pollination, nutrient cycling, food web dynamics and pest regulation, make them an important factor in agriculture, economy and health (Satterfield et al., 2020; Wotton et al., 2019). Migratory insects not only influence the ecosystems at their emergence site but also connect ecosystems across, sometimes vast, distances and over several life cycle stages and multiple generations (Darnis et al., 2025; Stefanescu et al., 2013; Suchan et al., 2024). Hoverflies (Syrphidae), for example, pollinate wild plants and crops while their larvae predate on aphids – both of which yield significant, yet hard-to-quantify economic benefits to agriculture (Reynolds et al., 2024). Conversely, migratory insect pests such as fall armyworm damage maize plants (e.g. Banson et al., 2020; Tao et al., 2022) or various plant- and leafhoppers play a central role in pathogen transmission to economically important crops (Nault and Ammar, 1989; Plante et al., 2024).

Insects have long been known to migrate over long distances (e.g. Hardy and Milne, 1938; Johnson, 1957; Taylor, 1974), and they are thought to be more flexible than migratory vertebrates in terms of routes and timing, and more dependent on environmental cues such as wind, temperature and precipitation. However, their small sizes and often high-altitude movements make it difficult to describe flyways or quantify the transport of biomass within or between regions (Chapman et al., 2015). Consequently, quantifications of the abundance of insect migration, their seasonally preferred directions of movement and taxonomic composition remain scarce (but see Hu et al., 2016; Huang et al., 2024; Werber et al., 2025), even more so for regions with complex topography or heterogenous landscapes, potentially acting as barriers.

Here, we characterize insect migration across Switzerland using a remote-sensing approach with vertical looking radars. Switzerland’s landscape is highly diverse, spanning the starkly different topographies of the Alps, Jura mountains, and pre-Alpine lowlands. Although topography is known to shape migratory movements of vertebrates (Beason, 1978; Bruderer and Jenni, 1990; Hirschhofer et al., 2024; Kranstauber et al., 2023), its role for those of insects has been studied only sporadically and exclusively using field-level methods that do not measure movements higher up in the air (Hawkes et al., 2024). We measured aerial insect movements from spring to autumn 2022 with vertical-looking radars placed at three topographically contrasting locations at 510 m asl in the pre-Alpine lowlands, at 1350 m in the Juran mountains, and at 2210 m at an Alpine high-mountain pass, characterising spatial and temporal activity patterns, flight directions, and community compositions.

## Results

### Insect activity

A total of 6.6 million individual insects were recorded across the three sites during the 8-month study period. This corresponds to an estimated number of 21 billion medium- to large-sized insects moving across Switzerland, amounting to a biomass of roughly 800 t (Fig. 1). Mean insect traffic per site varied between 2,000 and 6,000 insects per hour and km (Fig. 2a). There was little insect activity outside the study period, as indicated by the continuously operating radar at the Lowland site, which recorded only 1.4% of total activity before mid-March or after mid-November. Insect activity showed a first small peak with more than 1000 insects per hour and km during daytime both in the Lowland and at the Alpine Pass (Fig. 2b; measurements on the Jura Mountain started only in mid-April). The first significant nocturnal insect movements took place at the end of April and beginning of May on the Alpine pass (Fig. 2c). Activities then steadily increased at all sites until mid-June and reached peaks of more than 20,000 insects per hour and km at the Alpine pass and above the Jura mountains (Fig. 2b and 2c), whereas such peaks were more frequent at the Alpine pass (e.g. daytime in the beginning of July). Later in the year, daytime insect activity steadily declined to once more increase slightly in October. Nocturnal activity remained high throughout August and declined rapidly as of September.

**Figure 1.**
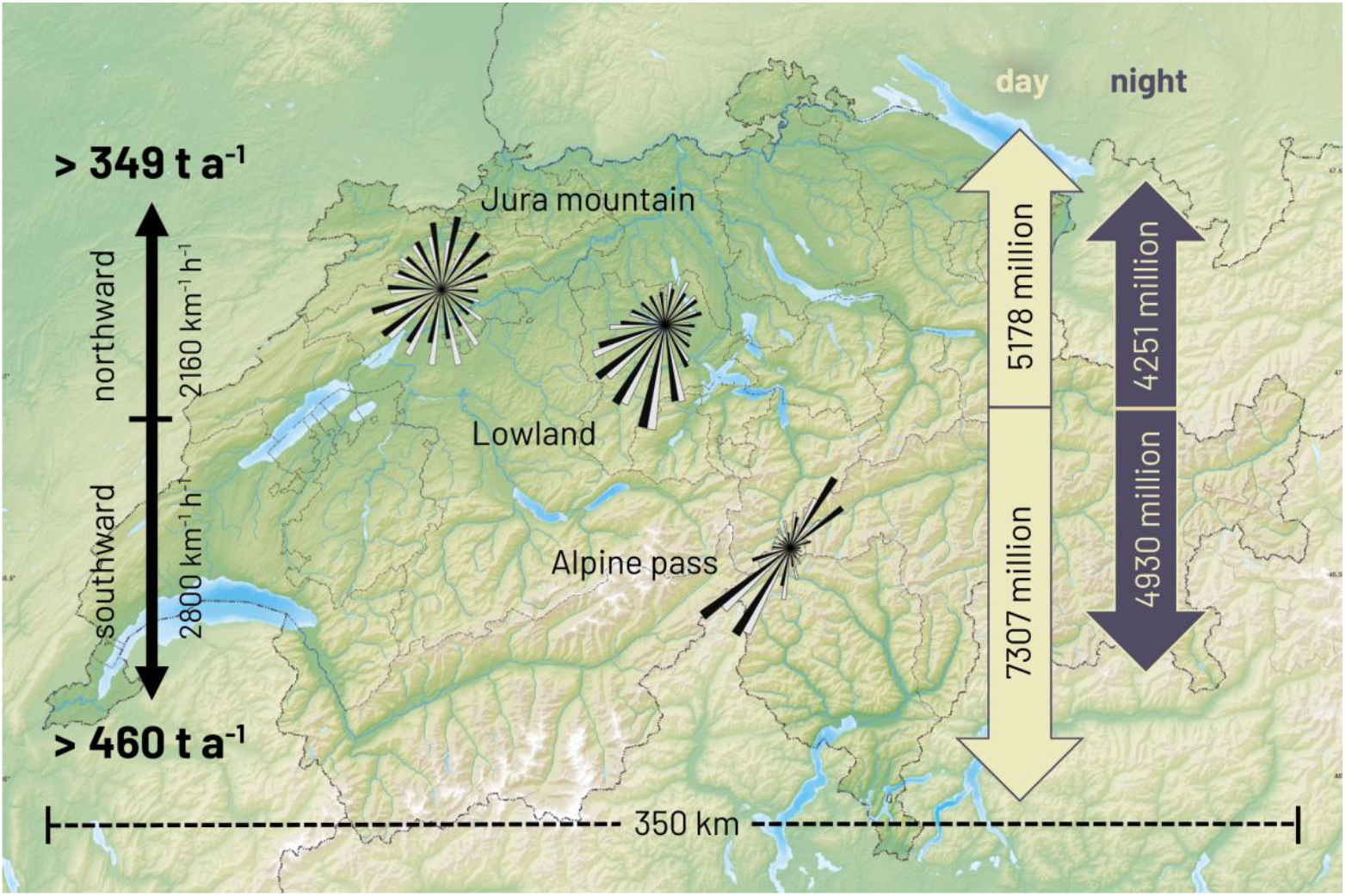
Measurement locations, movement directions, and total number of medium- to large-sized migratory insect movements and associated biomass flow across heterogenous terrain in Switzerland – an Alpine pass (southeastern location), the Jura mountains (western location) and in the Swiss Lowlands (middle). The total number of insects migrating during the day and night (colored arrows to the right) and the associated biomass crossing Switzerland to the north and south (black arrows to the left) were estimated as the sum over the entire observation period (15 March – 15 November 2022), extrapolated over the entire East-West extent of Switzerland (350 km). For each site, the circular plots indicate the relative frequency distributions of flight directions over day (white) and night (black). Topographic base map (c) Tschubby, CC BY-SA 3.0, https://commons.wikimedia.org/w/index.php?curid=17945004.

**Figure 2.**
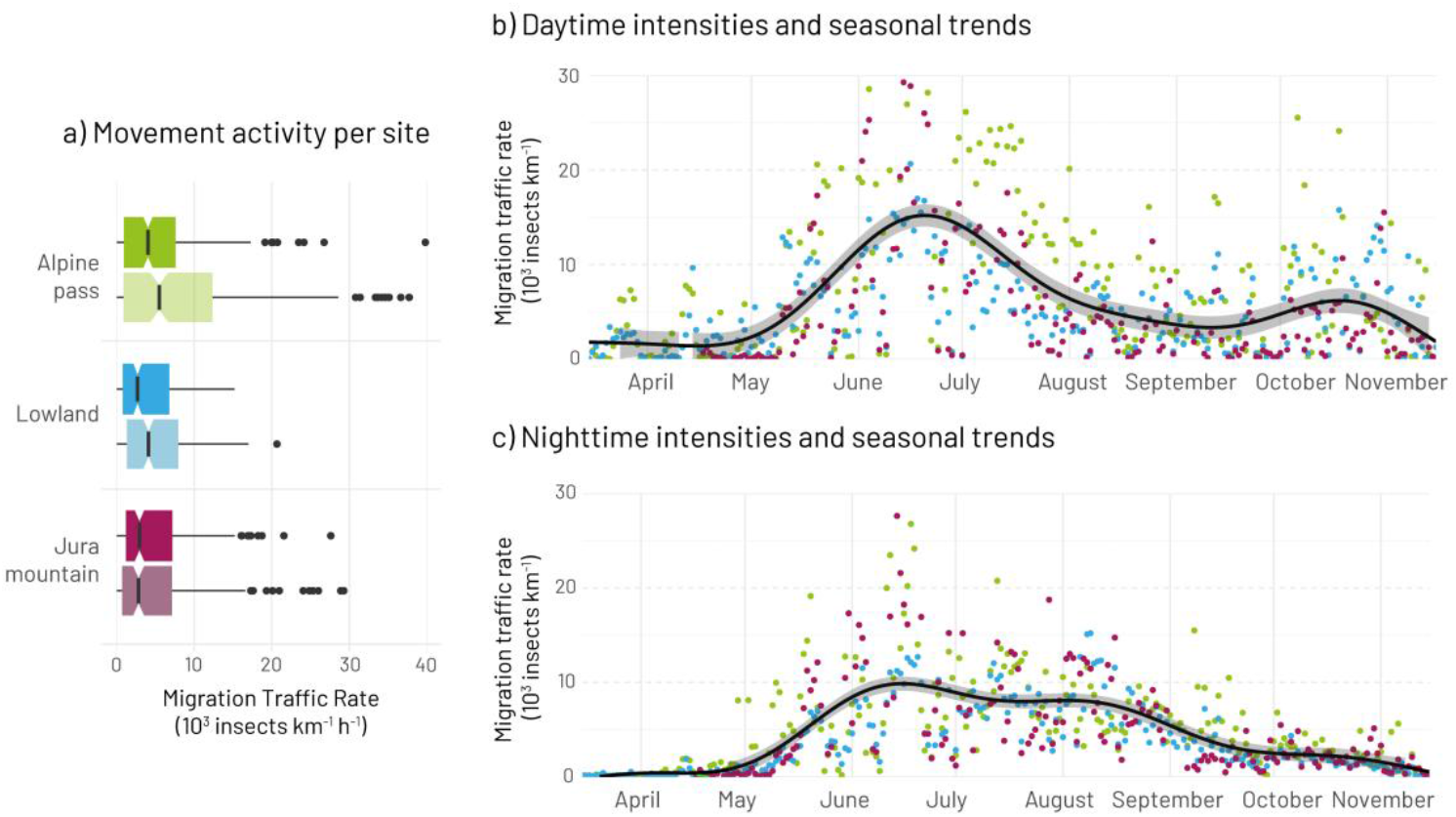
Insect traffic rates across the year at an Alpine pass site (green), the Lowlands (blue) and the Jura Mountains (purple); a) distribution of mean hourly insect traffic rates per site and day (light colours) and night (dark colours) - boxplots show median (center line), interquartile range (box), 95% CI of the median (notch), whiskers extending to 1.5×IQR, and outliers; b) mean **daily** insect traffic rates across the season (colored dots) and mean trend averaged across all sites (black line with 95% confidence range); c) mean **nightly** insect traffic rates across the season (colored dots) and mean trend averaged across all sites (black line with 95% confidence range).

After removing seasonal trends and site-specific autocorrelation, significant correlations remained between all three sites, both during day and night. The correlations were strong between Lowland and the Jura Mountain (day: 0.61; night: 0.46), moderate between Alpine pass and Lowland (0.44; 0.37), and weakest between Alpine pass and Jura Mountain (0.44; 0.25; for details, see suppl. Mat. Fig. S1 and S2).

Absolute insect activity, however, differed significantly between sites, with the Alpine Pass showing the highest overall activity despite lower temperatures (Fig. 2a and 3a; suppl. Mat 2.2). A large proportion of the variation in insect counts was explained by within-site temperature and variation between the sites and day versus night (Marginal R^2^ = 0.57 for fixed effects, Conditional R^2^ = 0.65 including random day-to-day variation; suppl. Mat 2.2). Nighttime activities were slightly lower than daytime activities (Fig. 3b). Within sites, insect abundance closely followed short-term, local temperature changes (Fig. 3a; also indicated by the strong positive effect in Fig 3b and suppl. Mat 2.2). Within-site temperature had the strongest influence on insect activity, followed by day/night effects and site differences (Fig. 3b).

**Figure 3.**
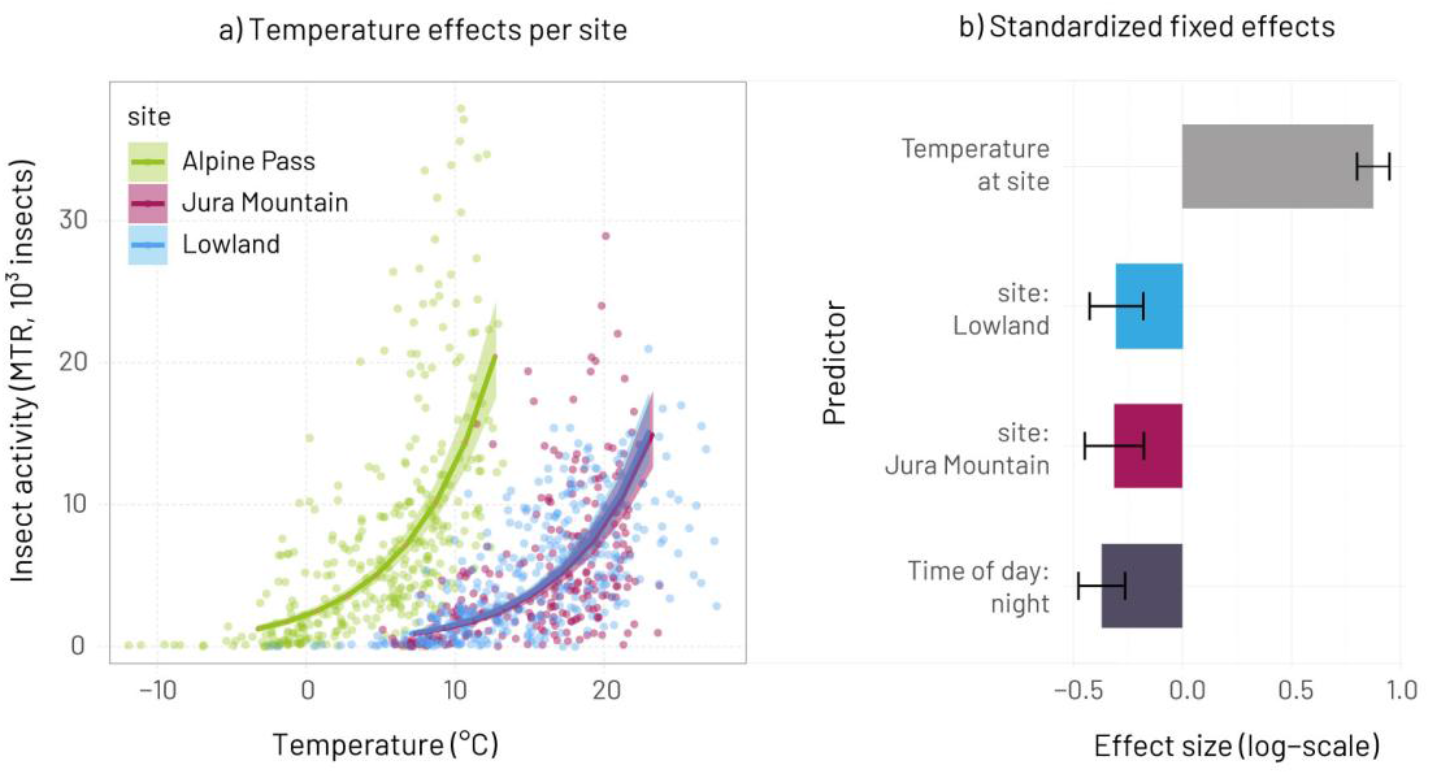
Result of the generalized linear mixed model (s. method and SI 2.2) showing the trend of activity in relation to ambient temperature at 100m agl (solid lines) with 95% confidence intervals (shades) and raw data (points). Colours correspond to the three sites.

### Flight directions

On the Alpine Pass, insects primarily flew along the valley axis, resulting in a clear northeast-southwest movement pattern (Fig. 1). A similar, albeit less pronounced, bi-directional pattern was observed at the Lowland site, whereas above the Jura Mountain, flight directions were highly scattered. The probability of southward flights differed strongly among sites and between day and night (Fig. 4; suppl. Mat 2.3). During daytime, the estimated overall mean proportion of southward flights across the study period was highest at the Lowland site (70%), followed by the Alpine pass (67%), and Jura Mountain site (65%). The nocturnal proportion of southward movements was similar to the daytime proportions at the Lowland site (67%) and Alpine pass (65%), but significantly lower at the Jura Mountain site (55%). Flight directions also changed over the season, and this seasonal change was different at each site. At the Alpine site, daytime southward directions increased steadily over the year. For the nocturnal movements, the increase in southern movements at the Alpine site was rather sudden with the switch towards more southward movements occurring between June and July. At the Lowland site, southward movements dominated throughout, but especially from June onwards. At the Jura Mountain site, daytime directional preferences hardly varied throughout the season. The proportion of nocturnal southward movements had a similar seasonal pattern to that of the Alpine site (Model fit: R^2^_adj_ = 0.12, deviance explained = 9.36%, for further details see suppl. Mat. 2.3).

**Figure 4.**
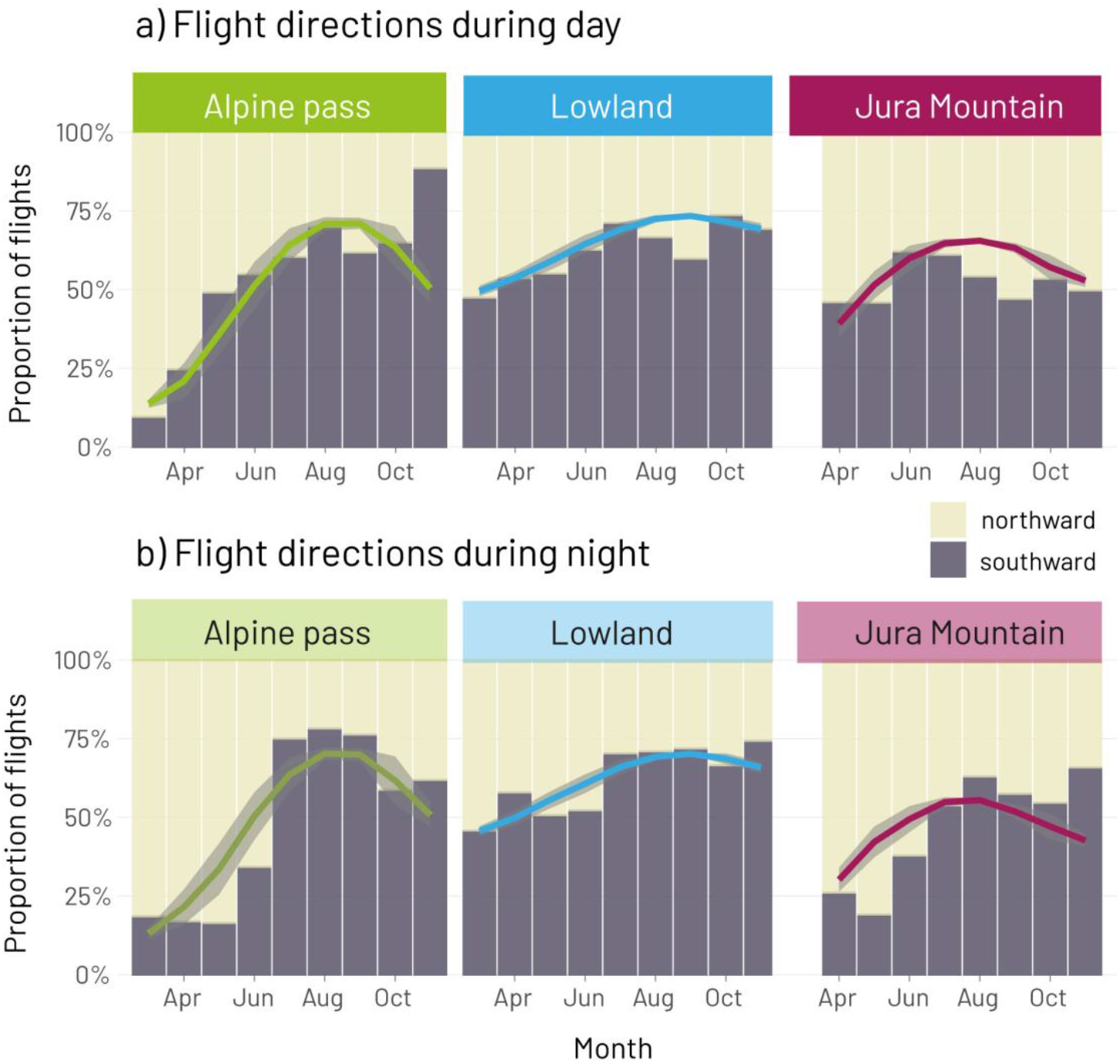
Monthly proportions of (a) daily and (b) nightly north- and southward insect movements for Alpine pass (left panel), Lowland (middle panel) and Jura mountain (right panel). Except for early in the season, southward directions dominated movements over the day at all sites. During nighttime, southward movements dominated in the Lowland site throughout the study period but only in the second half of the year for Alpine pass and Jura Mountain.

### Insect community composition

Wingbeat frequencies were significantly lower at night than during the day (night vs day: 35 vs 52 Hz; Fig. 5; suppl. Mat 2.4). Site was a significant predictor of wingbeat frequency; however, its effect size was small relative to unexplained within-group variability (residual SD = 13.3 Hz). The effect of time of day on wingbeat frequency differed among sites, with night–day differences being largest at the Jura mountain and smallest at the Alpine pass. (suppl. Mat. 2.4). Overall, the model explained only a minor proportion of the variation in the wingbeat frequencies (Marginal R^2^ = 0.15 for fixed effects, Conditional R^2^ = 0.18 including random month-to-month variation).

**Figure 5.**
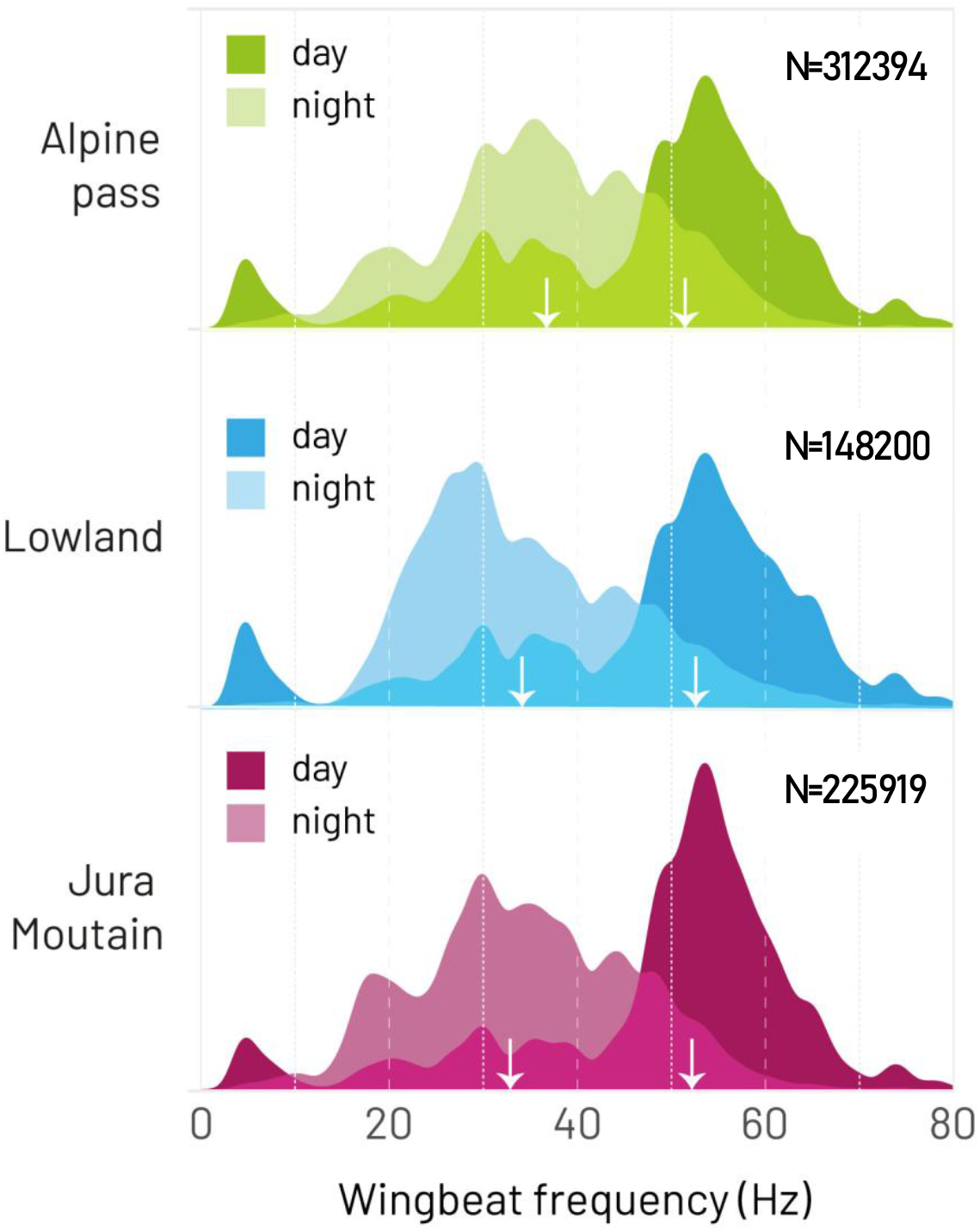
Relative distribution of wingbeat frequencies per site and time of day summarized across the whole observation period. Only wingbeat frequencies with a probability of fit >0.8 where included (see SI 2.5.). Sample sizes for day and night: Alpine pass N = 120,211/192,183, Lowland N = 48,650/99,185 and Jura Mountains N = 64,137/161,782. White arrows indicate medians for day and night, respectively.

## Discussion

We quantified insect biomass flows at contrasting sites within a complex mountainous landscape, characterizing seasonal dynamics, spatial coherence, predominant flight directions, and community composition. Our study yields several overarching insights: (a) Insect activity exhibited strong seasonal dynamics driven by local temperature, with activity patterns that were spatially coherent across topographically heterogeneous sites, resulting in an inferred minimum aerial insect biomass flow of ∼800 t per year across Switzerland; (b) the Alps do not appear to constitute a major barrier to migrating insects; (c) insects exhibit seasonally preferred flight directions; and (d) the composition of the aerial insect community differs markedly between day and night, but much less among sites.

Based on our measurement and extrapolation to Switzerland, we estimate an annual biomass flow of approximately 800 t of medium- and large-sized insects (corresponding to 2.3 t per km and year). However, our approach inherently underestimates total biomass because it does not detect small insects or individuals flying above the altitudinal measurement range of the radar. To place our estimates in context, we compare them with those of a comparable radar-based study by (Hu et al., 2016). Hu et al. (2016), while accounting for differences in detection sensitivity and altitude coverage, sampled insects between 150 and 1200 m above ground level, thereby missing low-flying insects but covering higher altitudes than our measurements. Accordingly, they detected approximately one third of insect activity above 500 m, whereas we observed about 50% of activity below 150 m. Despite these methodological differences, both studies yield estimates of the same order of magnitude (600–800 t of insect biomass per year). Applying a rough correction for undetected small insects (<10 mg), following Hu et al. (2016), suggests that total aerial insect biomass flow over Switzerland could be around 10,000 t per year.

Despite the importance of (aerial) insects for a wide range of ecosystem functions, relatively few studies have quantified their abundance or biomass fluxes. Existing estimates are based on a combination of ground-based trapping or visual observations (Grüebler et al., 2008; Hails, 1982; Hawkes et al., 2022; SHORTALL et al., 2009), and, more recently, remote-sensing approaches. For example, using a similar biological radar, the biomass of nocturnal insects in an agricultural landscape in eastern China was estimated to average 12.7 t per km and year, approximately an order of magnitude higher than our nocturnal estimate (Huang et al., 2024), suggesting that there are compelling differences between continents and/or habitats. Weather-radar–based approaches provide an alternative perspective at much larger spatial scales but rely on indirect inference of insect numbers from radar reflectivity (Dokter et al., 2019). Using weather radar data, Mungee et al. (2025) estimated mean insect densities of around 21 million insects per km^3^ at night and 46 million insects per km^3^ during the day across the United Kingdom. Although these values are not directly comparable to our measurements, converting them into traffic rates by assuming a flight speed of 5 km per hour yields insect traffic rates several orders of magnitude higher than those observed here. Notably, these estimates were derived from a relatively narrow altitudinal band (500–700 m above sea level), which may partly account for the high inferred densities and underscores the need for future confirmation.

Our results suggest that insects migrate heavily along mountain passes to cross the Alps, even under low air temperatures and at high altitudes. Various field-based studies have documented the funneling of migrating insects through narrow corridors when crossing mountain ranges (Aubert, 1964; David and Lack, 1951; Hawkes et al., 2024; Prell, 1925), similar to the well-established role of mountain passes for migratory birds (Aschwanden et al., 2019; Liechti et al., 1996; Williams et al., 2001). Although insect activity would generally be expected to decline at higher altitudes due to lower temperatures, the Alpine pass exhibited the highest overall insect traffic rates despite consistently colder conditions (Fig. 3). More than 50% of recorded flights at the Alpine pass occurred at air temperatures below 10 °C. This suggests that many of these movements were not initiated locally but instead represent migratory flights initiated elsewhere under warmer conditions that were sustained throughout areas of lower temperatures. The strongly aligned flight directions along the valley axis at the Alpine site further indicate that we captured a large-scale, but locally funnelled, flux of insects crossing the Alps rather than predominantly local, e.g., foraging movements.

An earlier study using the same radar systems as in our study but conducted over spatially more confined setting and a much shorter time period (Knop et al., 2023) reported similar average insect traffic rates and comparable differences between alpine and lowland sites. Along a similar line as for our Lowland and Jura Mountain sites (Fig 2 and 3), they did not find a difference between their lowland rural and urban landscape sites. Taken together, these findings suggest that insect flight activity over Switzerland is remarkably coherent across space, despite pronounced differences in elevation, terrain structure, and local climatic conditions. Although Alpine topography clearly influences flight activity patterns and movement directions (Fig. 3 and 4), the Alps overall do not appear to constitute a barrier to insect migration.

Preferred flight direction (northward vs southward) varied considerably over the study period, between sites, and between day and night (Fig. 4). A seasonal shift from predominantly northward to southward movements would be expected for migratory insects and was most pronounced at the Alpine pass, once more suggesting many movements at the Alpine pass are migratory. Preferred nocturnal flight directions were also similar between the Alpine pass and the Jura mountains, suggesting that large-scale migratory movements at higher altitudes extend across wide areas. In contrast, seasonal changes in flight directions were weaker at the Lowland site and also in general during daytime compared to nocturnal movements. Insects, thus, seem to have seasonally structured migratory directionality, although it is less sharply defined than in other aerial migrants such as birds (Tschanz et al., 2020).

The composition of the aerial insect community is known to differ considerably between day and night (Guevara and Avilés, 2013), with the nocturnal community being bigger on average (Haest et al., 2024), a pattern which is reflected in the lower nocturnal wingbeat frequency distributions observed in our data (Fig. 5). Wingbeat frequencies, however, differed little among sites; although these differences were statistically significant, they had small effect sizes. The single exception was the nocturnal insect community at the Alpine pass, where wingbeat frequencies were higher on average than at the other sites. This pattern contrasts with the expectation that larger-bodied insects, which typically have lower wingbeat frequencies, should be better able to withstand low nighttime temperatures. Previous light-trap studies have indeed reported higher moth biomass at higher elevations, and a greater contribution of larger insects in the Alps compared to the Swiss lowlands (Fontana et al., 2020). However, in birds, wingbeat frequency increases with altitude due to reduced air density (Pennycuick, 1996; Schmaljohann and Liechti, 2009). Whether similar aerodynamic constraints influence insect flight at high altitudes remains unclear (Dillon et al., 2006; Dudley, 2000) and warrants future studies.

Clearly, our study is only a first step in unravelling the magnitude and extent of continental insect biomass flows. For context, nocturnal bird migration over Switzerland has been estimated at approximately 230 million individuals per year (Nussbaumer et al., 2021); which, assuming a mean body mass of 30 g, corresponds to roughly 7,000 t of avian biomass annually. Against this backdrop, the estimated insect biomass flow of about 10,000 t per year is substantial and appears to exceed that of migratory birds. A comprehensive assessment of continental insect biomass flows and their consequences for agriculture, ecosystem functioning, and the spread of diseases, will require sustained, continuous remote-sensing observations across multiple sites that span broad climatic, biome, and habitat gradients, and that extend over multiple years.

## Materials and Methods

### Data collection

We recorded aerial movements of individual insects with BirdScan MR1 X-band radars (https://swiss-birdradar.com/) at three sites in Switzerland (Fig. 1). The three sites lay along a northwest–southeast transect of about 100 km, spaced at intervals of roughly 50 km. The eastern site was an Alpine pass at 2210 m asl (‘Alpine pass’, 46.552337°N, 8.763809°E), which runs from north to south and is surrounded by mountains of up to 3100 m asl. The middle site lay to the northwest of the alpine site, in the central lowlands at 510 m asl (‘Lowland’, 46.533811°N, 8.587433°E). The site is surrounded by rolling hills of up to 800 m asl and a small lake nearby. The western site (‘Jura Mountain’, 47.23194°N, 8.39694°E) was at 1350 m asl on the southernmost ridge of the Jura mountains. The site is surrounded by mountains of similar or lower heights to the North and West and a steep slope to the south rises from 430 m to 1330 m asl, marking the northern border of the Swiss lowlands. For the Alpine and Lowland site continuous measurements were made from 15 March to 16 November 2022, for the Jura Mountain site from 17 April to 16 November 2022.

The vertical looking radar detects targets from about 40m upwards. Individual insects >=0.1 g can be detected at least up to 500 m, while very small insects of <0.01 g are hardly detected at all. For all analyses we only included insect targets recorded between 40 and 500m (n = 6,613,498). For the flight direction analysis, we only retained individuals for which the observed flight path was at least half of the aerial diameter of the radar beam at the respective altitude. Using these 41% of individuals with such longer flight tracks (n = 2,704,414) excludes insects that only briefly crossed the radar beam and whose flight directions can thus not be reliably estimated.

We analysed wingbeat frequency distributions to compare the community composition of insects between the sites. The wingbeat frequency of each insect was estimated using a random forest regression based on a labelled dataset (Haest et al., 2021). Only insects with a wing beat frequency probability of fit of higher than 0.8 were included - a threshold that was set empirically by comparing frequency distributions resulting from different threshold values (0.5 – 0.9). Using a 0.8 threshold yielded a sufficiently robust wingbeat frequency distribution and a sample size for the analysis of the insect community composition that remained representative of the overall community composition (n_tot_ = 686,513; for each site between 9 and 13% of all samples; see suppl. Mat. Fig. S4). Further details of the radar system are provided in Zaugg et al. (2008) and Schmid et al. (2019), whereas the measurement scheme is described in Haest et al.(2024).

### Analysis

We investigated variation in flight activity, flight directions, and the composition of wingbeat frequencies between the sites, and how activity and directional patterns corresponded across sites over time. Detections of individual insects were aggregated into non-directional insect traffic rates, a standardized aerial flight activity measure that quantifies how many insects per hour cross a virtual line of one km and makes measurements comparable between radar systems (Bauer et al., 2024; Haest et al., 2024; Schmid et al., 2019). Calculations were done with the *‘birdscanR*’ R-package (Haest et al., 2025) and resulted in a time series of hourly values of insect traffic rates for each of the three sites.

To remove shared seasonal patterns and temporal autocorrelation, we fitted a generalized additive mixed model (GAMM) using the *mgcv* package in R. Insect traffic rate was modelled as a smooth function of day of year, with site included as a random intercept. Temporal autocorrelation within sites was accounted for using a first-order autoregressive [AR(1)] correlation structure. Normalized residuals from the fitted model were subsequently extracted and used to assess whether residual variation remained correlated between sites, indicating spatial synchrony beyond shared seasonal dynamics (Spearman, pairwise correlation).

We estimated the overall insect biomass flow over Switzerland by calculating the mean hourly insect traffic rate for all sites across the entire measurement period (15 March 2022 00:00 – 16 November 2022 00:00) and multiplying it by the total number of hours, the maximum geographic East-West range of Switzerland (350 km), and a mean insect body mass of 38 mg (based on Hu et al. (2016); for details: see suppl. Material table S1 and S2).

To investigate variation in insect activity due to site, daytime, and temperature, we fitted a generalized linear mixed model using a negative binomial error distribution. Site, within-site centered temperature, and day/night were included as fixed effects, and day of year was included as a random intercept to account for daily variability. We used within-site centered temperatures to separate short-term temperature effects from between-site differences (R-package ‘*glmmTMB’*). We downloaded hourly temperature data for pressure levels at approximately 100 m above ground level for each site (Alp = 750 hPa, Low = 925 hPa, Jur = 850 hPa) from the ERA5 reanalysis data (Muñoz Sabater, 2019), with a spatial resolution of around 9 km (0.1° x 0.1°).

Flight directions were analysed as a binary response variable. To assess how the probability of southward flight varied among sites, between day and night, and over the course of the year, we fitted a generalized additive model (GAM) with a binomial error distribution and logit link function, using the ‘*bam’* function from the R-package *‘mgcv’*. The model included diel period (day, night) and their interaction as parametric fixed effects. Seasonal patterns were modeled using smooth functions of day of year. As seasonal dynamics were expected to differ among sites, the model included both a global smooth of day of year and site-specific smooths (suppl. Mat. 2.3). Day-to-day dependence and unmeasured short-term influences (e.g. weather conditions) were accounted for by including calendar date as a random effect. Spline basis dimensions (k) were chosen conservatively, and final smoothness was determined by the automatic penalization implemented in *mgcv*, reducing the risk of overfitting. Given the large sample size (>2.7 million observations), models were fitted using the discrete approximation and estimated via fast restricted maximum likelihood.

Differences in the insect community composition were inspected by analysing the distribution of wingbeat frequencies between sites and daytime using a linear mixed-effects model with site, day/night, and their interaction as fixed effects, and month included as a random intercept (R-package ‘*lmer4’* (Bates et al., 2015)). For all analyses, we used R version 4.5.1 (R Core Team, 2025).

If your research involved human or animal participants, please identify the institutional review board and/or licensing committee that approved the experiments. Please also include a brief description of your informed consent procure if your experiments involved human participants.

## Acknowledgments

We thank the energy supplier “Stadtwerke Grenchen” (SWG; Jura Mountain) and the Federal Office for Armaments Procurement (armasuisse; Alpine pass) for funding the data collection and making it publicly accessible. Fabian Hertner from Swiss BirdRadar Solution AG gave assisted with radar data access and processing. This project is part of HiRAD (https://hirad.science), funded through the 2022–2023 Biodiversa+ BiodivMon call for research proposals, with the funding organizations Swiss National Science Foundation, Switzerland (SNF 31BD30_216840), Belgian Federal Science Policy Office, Belgium (BelSPO RT/24/HiRAD), Netherlands Organization for Scientific Research (NWO EP.1512.22.003), and Academy of Finland, Finland aka 359864).

