## Supplemental information for "A hidden biomass flow - 10,000 tons of insects fly over Switzerland"

1. From numbers to biomass - refer to “insect activity”

To convert the number of insects into a biomass number (tons), we had to define an average weight for a single individual. Similar to the radar used in the study of (Hu et al., 2016), our radar records medium to large sized insects. We therefore used the same proportions of medium and large sized insect to calculate a mean insect body mass for our sample (Table S1). We divided their estimated biomass for medium sized insects through the number of medium sized individuals and the estimated biomass for large sized insects through the number of large sized individuals to achieve an average body mass for medium and large insects (26mg / 146mg; J. Chapman pers. com.).

**Table S1: Calculating a mean insect body mass for our sample based on the figures taken from (Hu et al., 2016).**

| **Size** | **N [10^6]** | **%** | **Total [t]** | **Mean [g]** |
|---|---|---|---|---|
| medium | 14650000000 | 0.90 | 380.8 | 0.026 |
| large | 1560000000 | 0.10 | 227.59 | 0.15 |
| *total/mean* | *16210000000* | *1* | *608.39* | *0.038* |

Finally, we used the weighted average of 38 mg, based on the proportions (0.9/0.1) given by (Hu et al., 2016) for the two groups, and calculated the corresponding biomass for Switzerland (Table S2).

**Table S2: Calculating the insect biomass including all individuals (medium and large insects) recorded in this study.**

| direction | No/(year*km) | mean weight | range [km] | CH mass [t/year] |
|---|---|---|---|---|
| southward | 35000000 | 0.038 | 350 | 460 |
| northward | 26600000 | 0.038 | 350 | 349 |
| total | 61600000 | | | 809 |

2. Statistical analysis

2.1. Temporal correlation between sites – refer to "insect activity"

**Figure S1:** Pairwise correlation for the site specific residuals of insect traffic rates in relation to the overall seasonal **day** trend. Shown are raw residuals with regression lines and 95% confidence limits, density distributions, Pearson's ρ-values, and sample sizes.

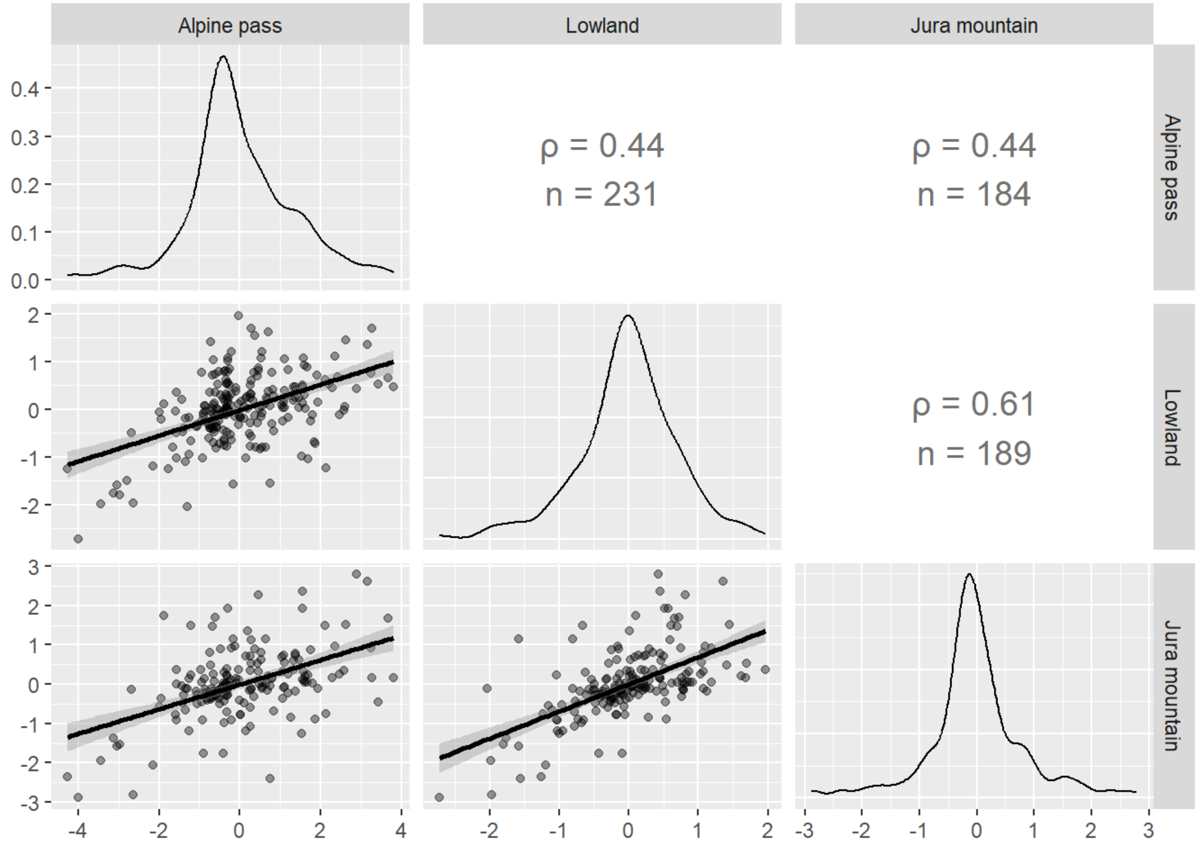

**Figure S2:** Pairwise correlation for the site specific residuals of insect traffic rates in relation to the overall seasonal **night** trend. Shown are raw residuals with regression lines and 95% confidence limits, density distributions, Pearson's ρ-values, and sample sizes.

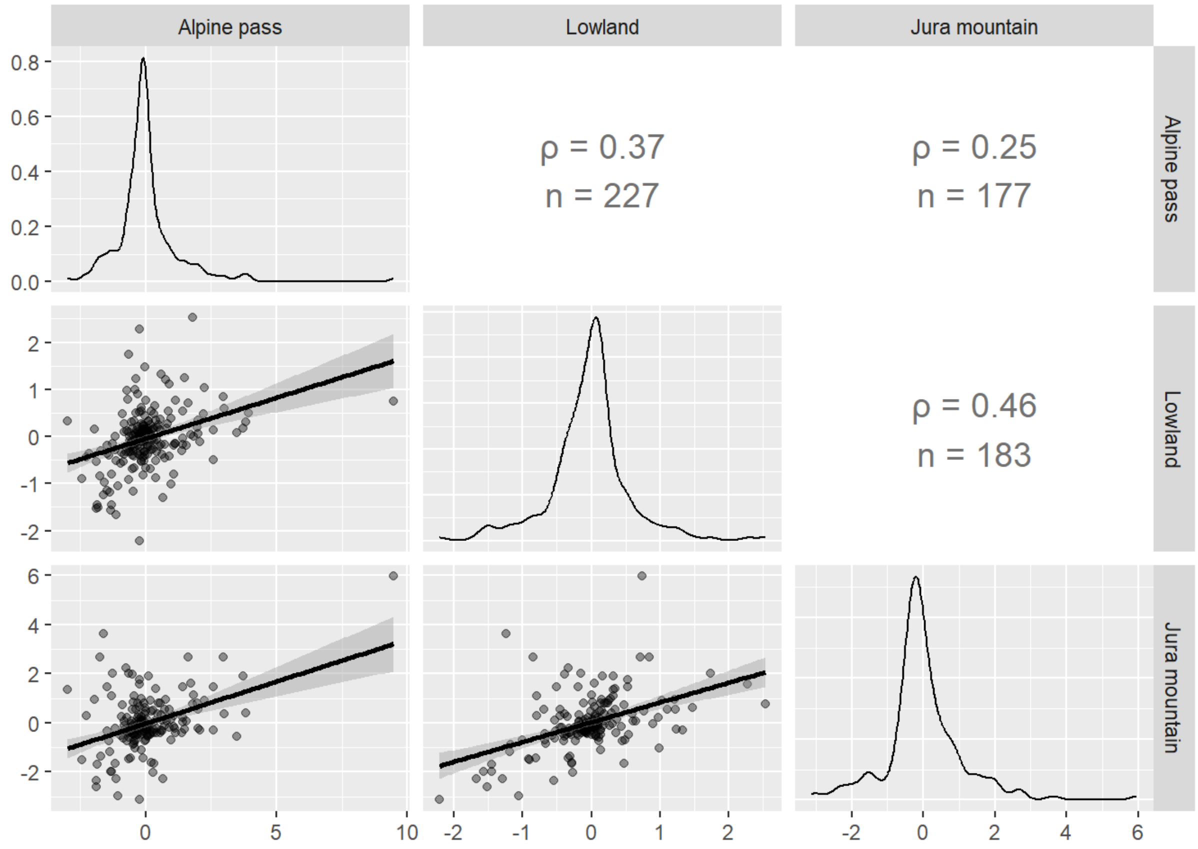

##### *2.2.* Modelling Insect traffic rate (ITR)

**Model structure:**
glmmTMB( ITR ~ site + Temperature_within_site + dayOrNight + (1 | dayofyear),
family = nbinom2)

**Site effects:**

| site | response | SE | df | asymp.LCL | asymp.UCL |
|---|---|---|---|---|---|
| Alpine Pass | 5044 | 261 | Inf | 4558 | 5582 |
| Jura Mountain | 3691 | 219 | Inf | 3287 | 4145 |
| Lowland | 3728 | 189 | Inf | 3374 | 4118 |

*Results are averaged over the levels of: dayOrNight*
*Confidence level used: 0.95*
*Intervals are back-transformed from the log scale*

**Temperature effects within sites:**

| Temp within site | response | SE | df | asymp.LCL | asymp.UCL |
|---|---|---|---|---|---|
| -2 | 2895 | 115 | Inf | 2677 | 3130 |
| 0 | 4109 | 155 | Inf | 3817 | 4424 |
| 2 | 5834 | 241 | Inf | 5380 | 6326 |

*Results are averaged over the levels of: site, dayOrNight*
*Confidence level used: 0.95*
*Intervals are back-transformed from the log scale*

**Standardized parameters:**

| Parameter | Std_Coefficient | CI | CI_low | CI_high |
|---|---|---|---|---|
| (Intercept) | 8.7106690 | 0.95 | 8.5939819 | 8.8273561 |
| siteJura Mountain | -0.3122838 | 0.95 | -0.4479474 | -0.1766201 |
| siteLowland | -0.3023895 | 0.95 | -0.4250775 | -0.1797015 |
| Temp_within_site | 0.8759800 | 0.95 | 0.8016667 | 0.9502932 |
| dayOrNightnight | -0.3695465 | 0.95 | -0.4763776 | -0.2627153 |

**R2 for Mixed Models**
Conditional R2: 0.654
Marginal R2: 0.569

2.3. Modelling directional preferences in space and time

**Model structure:**

```
direction_bin ~ site * day_night + s(doy, k = 8) + s(doy, by = site, k = 4) + s(date_f, bs = "re")

Parametric coefficients:
                                    Estimate     Std. Error    z value    Pr(>|z|)
(Intercept)                         -0.606729    0.041849      -14.50     <2e-16 ***
siteJura Mountain                   0.268143     0.024942       10.75     <2e-16 ***
siteLowland                         1.092889     0.012238       89.31     <2e-16 ***
day_nightnight                      -0.048504     0.004292     -11.30     <2e-16 ***
siteJura Mountain:day_nightnight    -0.375508    0.006777      -55.41     <2e-16 ***
siteLowland:day_nightnight          -0.110868    0.006914      -16.04     <2e-16 ***
---
Signif. codes:  0 '***' 0.001 '**' 0.01 '*' 0.05 '.' 0.1 ' ' 1

Approximate significance of smooth terms:
                             edf          Ref.df       Chi.sq       p-value
s(doy)                       4.895e-03    5.246e-03    3.000e-03    0.956
s(doy):siteAlpine pass       2.981e+00    2.984e+00    7.772e+02    <2e-16 ***
s(doy):siteJura Mountain     2.949e+00    2.962e+00    3.127e+02    <2e-16 ***
s(doy):siteLowland           2.854e+00    2.876e+00    1.085e+02    <2e-16 ***
s(date_f)                    3.015e+02    3.530e+02    1.652e+05    <2e-16 ***
---
Signif. codes:  0 '***' 0.001 '**' 0.01 '*' 0.05 '.' 0.1 ' ' 1

Rank: 376/377
R-sq.(adj) =  0.122   Deviance explained = 9.36%
fREML = 4.1624e+06  Scale est. = 1        n = 2714832
```

**site-day effects:**

| site | day_night | prob | SE | df | lower.CL | upper.CL |
|---|---|---|---|---|---|---|
| Alpine pass | day | 0.665 | 0.0131 | 2714516 | 0.638 | 0.690 |
| Jura Mountain | day | 0.654 | 0.0134 | 2714516 | 0.627 | 0.680 |
| Lowland | day | 0.701 | 0.0124 | 2714516 | 0.677 | 0.725 |
| Alpine pass | night | 0.654 | 0.0134 | 2714516 | 0.627 | 0.679 |
| Jura Mountain | night | 0.553 | 0.0146 | 2714516 | 0.524 | 0.582 |
| Lowland | night | 0.667 | 0.0131 | 2714516 | 0.641 | 0.692 |

Confidence level used: 0.95

Intervals are back-transformed from the logit scale

*2.4.* Community composition (wingbeat frequencies)

**Model structure:**

lmer(WFF_predicted ~ site * dayOrNight + (1|month)

Linear mixed model fit by REML ['lmerMod']
WFF_predicted ~ site * dayOrNight + (1 | month)

REML criterion at convergence: 5497985

Scaled residuals:

| Min | 1Q | Median | 3Q | Max |
|---|---|---|---|---|
| -3.7715 | -0.5709 | 0.0771 | 0.6542 | 7.8408 |

Random effects:

| Groups | Name | Variance | Std.Dev. |
|---|---|---|---|
| month | (Intercept) | 6.811 | 2.61 |
| Residual | | 176.767 | 13.30 |

Number of obs: 686148, groups: month, 9

Fixed effects:

| | Estimate | Std. Error | t value |
|---|---|---|---|
| (Intercept) | 48.48334 | 0.87214 | 55.59 |
| siteLowland | -0.96569 | 0.08059 | -11.98 |
| siteAlpine pass | -1.48907 | 0.06529 | -22.81 |
| dayOrNightnight | -14.21977 | 0.06312 | -225.29 |
| siteLowland:dayOrNightnight | 1.61001 | 0.09678 | 16.64 |
| siteAlpine pass:dayOrNightnight | 4.45624 | 0.07963 | 55.96 |

Correlation of Fixed Effects:

| | (Intr) | stLwln | stAlpp | dyOrNg | stL:ON |
|---|---|---|---|---|---|
| siteLowland | -0.043 | | | | |
| siteAlpnpss | -0.051 | 0.531 | | | |
| dyOrNghtngh | -0.052 | 0.547 | 0.675 | | |
| stLwlnd:dON | 0.036 | -0.831 | -0.440 | -0.638 | |
| stApss:dyON | 0.040 | -0.429 | -0.815 | -0.782 | 0.503 |

Analysis of Variance Table

| | npar | Sum Sq | Mean Sq | F value |
|---|---|---|---|---|
| site | 2 | 899468 | 449734 | 2544.2 |
| dayOrNight | 1 | 19703503 | 19703503 | 111466.0 |
| site:dayOrNight | 2 | 584902 | 292451 | 1654.4 |

### R2 for Mixed Models

Conditional R2: 0.183
Marginal R2: 0.152

**Figure S3 -** refers to Figure 5. Estimated marginal means (± 95% CI) of wingbeat frequencies across sites during daytime and nighttime. Differences among sites were pronounced during the night but reduced at daytime.

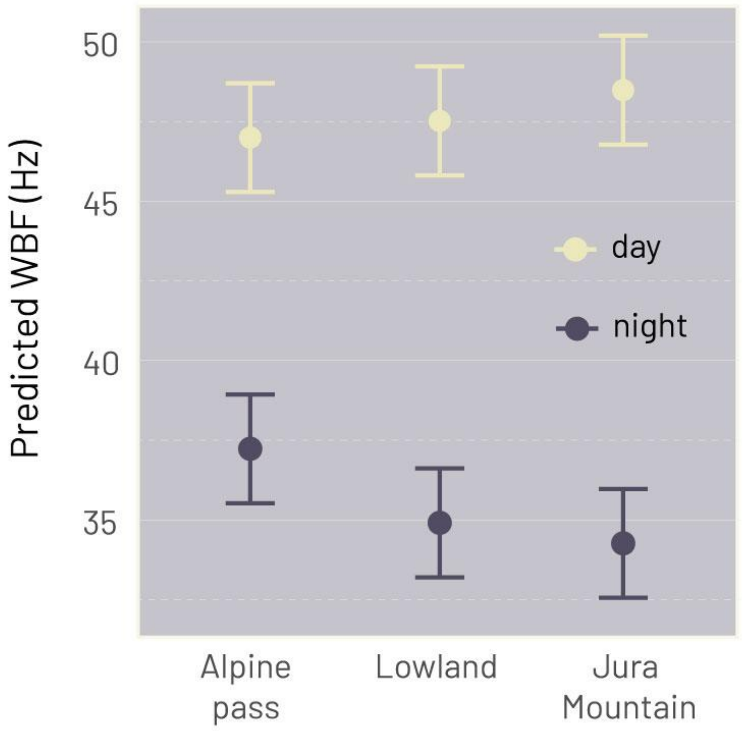

#### *2.5.* Threshold for wingbeat frequencies filtering

Wingbeat frequencies are extracted from the radar signal using a random forest classifier, based on various features of the signal (Zaugg et al., 2017, 2008). Aside from the predicted wingbeat frequency a probability of fit is provided. To decide which limits to be used for our analysis above, we plotted the wingbeat frequency distributions for various probability levels (see Fig S3). We decided empirically to choose probability level 0.8, as a compromise between accuracy and sample size.

**Figure S4:** Distribution of wingbeat frequencies for six probability levels (0.5 – 0.95). The plot with a probability level of 0.8 was selected for the study (different colours).

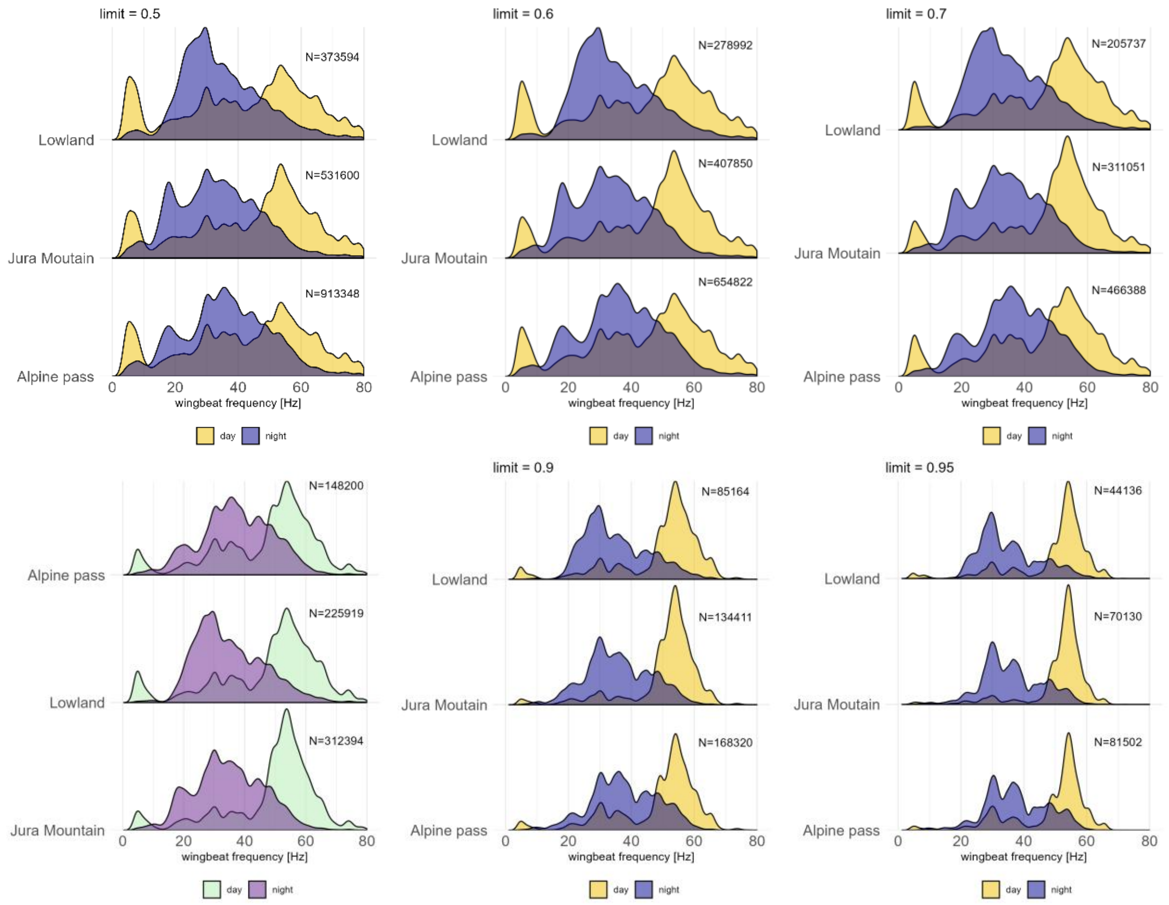
